# Post-sampling degradation of viral RNA in wastewater impacts the quality of PCR-based concentration estimates

**DOI:** 10.1101/2025.06.06.658243

**Authors:** James D. Munday, Jolinda de Korne-Elenbaas, Charles Gan, Adrian Lison, Julien Riou, Christoph Ort, Timothy R. Julian, Tanja Stadler

**Affiliations:** Department of Biosystems Science and Engineering, ETH Zuürich, Switzerland; SIB Swiss Institute of Bioinformatics, Switzerland; Eawag, Swiss Federal Institute of Aquatic Science and Technology, Switzerland, Switzerland; Department of Epidemiology and Health Systems, Unisanté, Center for Primary Care and Public Health & University of Lausanne, Switzerland

**Keywords:** Wastewater, Pathogen surveillance, Viral degradation

## Abstract

Successful wastewater-based infectious disease surveillance programs depend on regular, reliable molecular detection of nucleic acids in municipal wastewater systems. This process is challenged by the gradual degradation of the viral content of the wastewater over time. Testing protocols are complex and often cannot be performed on site, resulting in delays between collection and testing. The evidence of continued degradation of viral RNA when stored at low temperatures is currently limited to a small number of studies with mixed results. Taking advantage of variable delays between sample collection and processing, we used a Bayesian temporal model and data from two winter periods of a national surveillance program in Switzerland to determine the rate at which the measured viral concentrations of four respiratory viruses declined as a result of RNA degradation between sample collection and processing. We found evidence of substantial degradation between the collection and processing of samples with daily rates of up to −0.28 (−0.38 − −0.19 95% CrI). We established that reduction in viral concentrations resulting from post-sampling degradation was responsible for a number of measurements falling below quantifiable limits. For one treatment plant, we estimate that 39 measurements fell below the limit of detection due to RNA degradation over the course of a single season. Measurements are more likely to be lost early in the seasonal epidemic when concentrations are still low. This delays consistent reliable measurement and sets back epidemiological assessments relevant to public health management strategies.

## 1 Introduction

Monitoring municipal wastewater systems for the presence of pathogens has gained interest in recent years, following the demonstration of successful monitoring of SARS-CoV-2 [1]. By 2023, SARS-CoV-2 wastewater monitoring programs had already existed in more than 43 countries worldwide [2]. These programs offer public health authorities an alternative to clinical data to track infection prevalence [1, 3, 4], variant emergence, and the detection of cryptic lineages in real time [5–8]. Following the success of the SARS-CoV-2 monitoring, there have been renewed efforts to monitor other respiratory pathogens in wastewater, including influenza A virus, influenza B virus, and Respiratory Syncytial Virus (RSV) [9–14].

Wastewater-based pathogen surveillance relies on the regular collection of sewage samples from a consistent set of sampling sites, which are then tested for the presence of the pathogen of interest. Quantitative Polymerase Chain Reaction (qPCR) or digital PCR (dPCR) testing can be used to estimate the concentration of virus in the wastewater [15, 16]. Fluctuations in measured viral RNA concentrations in the wastewater can be used as a proxy for variation in prevalence of infection in the population [2, 17, 18]. Wastewater-based monitoring offers some benefits over clinical surveillance or other active surveillance approaches. Primarily, the viral concentrations offer non-intrusive and consistent measurement of virus shed from well-defined geographical areas. Samples are available in real time and in principle with shorter delays than those of clinical case reports and hospitalisations [5, 17]. Wastewater is also not sensitive to variation in reporting due to changes in healthcare-seeking behaviour and government-prescribed testing protocols, as clinical data is [19–23].

There are, however, some important challenges. Besides changes in viral concentration due to varying population prevalence of infection, there are a number of factors that contribute to variation in viral concentration in wastewater. For example, shedding distributions can vary between individuals [24–26]. The daily volume of influent wastewater at a sampling site can vary depending on multiple factors, including the topology of the wastewater system, the variation in the production of residential and industrial waste, and rainfall, which can cause the dilution of virons shed by infected individuals to vary from day to day. Another source of variability in viral concentration arises due to the variable degradation of RNA in wastewater [27]. After being shed from the body, RNA strands begin to break down resulting in fragmented RNA. Molecular methods like dPCR are used to quantify pathogens in wastewater by detecting pathogen-specific segments on this fragmented RNA. However, if fragmentation occurred within the region of the genome targeted by PCR, the signal is disrupted, leading to lower concentration estimates. The degradation rate of RNA is known to vary depending on environmental conditions. For example, laboratory-based analyses have shown variability in viral degradation with temperature [28] and the concentrations of various substances in the water [29]. However, it is not clear how these results translate to wastewater systems or how degradation rates could vary between the pathogens we monitor in a real-world context.

Samples can be stored in refrigerated (4^*o*^C) conditions for several days prior to testing, although degradation reduces at low temperatures, meaningful reduction in viral concentration has been measured in previous laboratory-based studies [28, 30, 31]. It is not clear how degradation in the interval between sample collection and processing may further affect the measured concentration. Further fragmentation of RNA can affect the quality of the resulting data and the epidemiological inferences that can be made.

As part of our routine monitoring of viral concentrations, we used dPCR to quantify concentrations of SARS-CoV-2, influenza A and B and RSV RNA in wastewater collected at six wastewater treatment plants (WWTPs) in Switzerland over a period of two years [14]. The samples were collected at each WWTP between 5 and 7 days a week. The samples were processed once or twice per week during this time. The varying delay between collection and testing provides a natural experiment to quantify degradation of RNA for each pathogen we collect, and in each WWTP we monitor. We noticed periodic patterns in the variation of viral concentrations we estimated. In several time series, it was clear that viral concentrations dipped most in measurements where the delay between collection and testing was greatest (Supplementary Material Figure 3). To evaluate these patterns further, we constructed a statistical model to quantify the relationship between testing delay and viral concentrations in wastewater while accounting for temporal correlation, time-varying population size and daily flow rates. Using our model, we quantified the impact of delays between collection and testing on the quality of our viral concentration measurements.

## 2 Methods

### 2.1 Wastewater data

Twenty-four hour composite wastewater samples were collected from six WWTPs across Switzerland five to seven days per week (Figure 1). Following collection, they were stored on site at 4^*o*^C for up to a week. We chose to analysed data from two winter seasons (October to March) as viral concentrations of influenza and RSV were low during the summer periods, making reliable concentriation quantification impossible. In July 2023 the processing protocol changed, therefore the two seasons followed different testing regimes. During the season between October 2022 and March 2023, batches of samples were transported on ice to the laboratory twice a week and processed directly after arrival. The laboratory had two processing regimes in place that aimed to minimize the time between collection and processing for all samples. In both regimes, samples from Monday and Tuesday were processed on Thursday of the same week. Samples from WWTPs in Altenrhein, Chur and Sensetal followed Regime A, with samples from Wednesday to Sunday being processed on the following Tuesday.

**Fig. 1.**
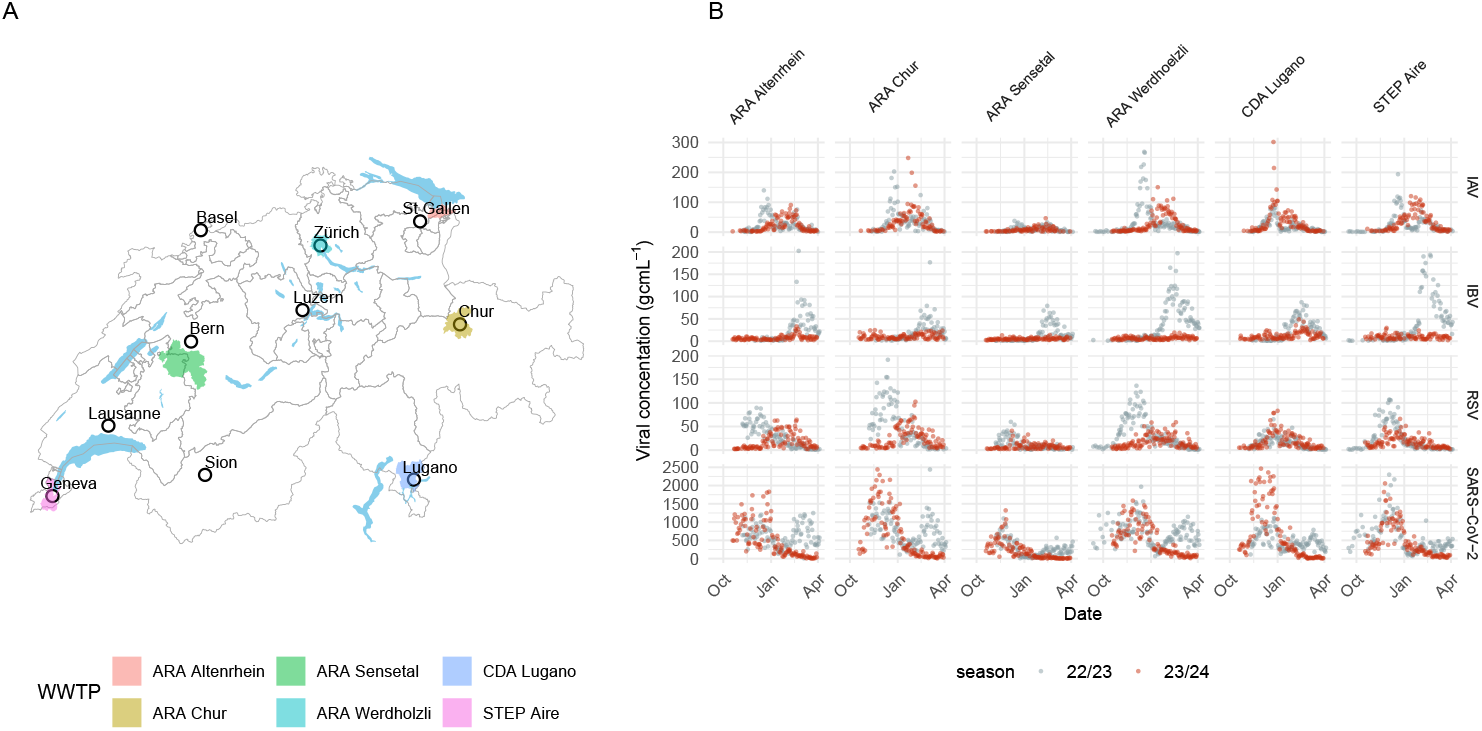
A)The location of treatment plants included in this study. Coloured polygons show the geographical extent of each treatment plant’s catchment. B) Viral concentrations of influenza A (IAV), influenza B (IBV), Respiratory Syncytial Virus (RSV) and SARS-CoV-2 (panels vertical) in each wastewater treatment plant (panels horezontal) over winter seasons in 2022-23 (grey) and 2023-24. (red)

Samples from WWTPs in Werdhölzli, Lugano and Aire followed Regime B, with samples from Wednesday to Sunday being processed on the following Wednesday. In the season from October 2023 - March 2024 samples were processed once per week on one of three processing days. Samples from Werdhölzli and Chur were processed on Tuesdays, samples from Lugano, Sensetal and Altenrhein were processed on Wednesdays and samples from Aire (Geneva) were processed on Thursdays (Figure 2). The samples were processed by concentrating and extracting total nucleic acids using a column-based direct capture method (Promega Wizard® Enviro Total Nucleic Acid Kit). The concentrations of SARS-CoV-2, IAV, IBV and RSV RNA were quantified in the wastewater extracts by dPCR. For each measurement date, each sample was processed in duplicate; we used the mean of the two concentration estimates [14].

**Fig. 2.**
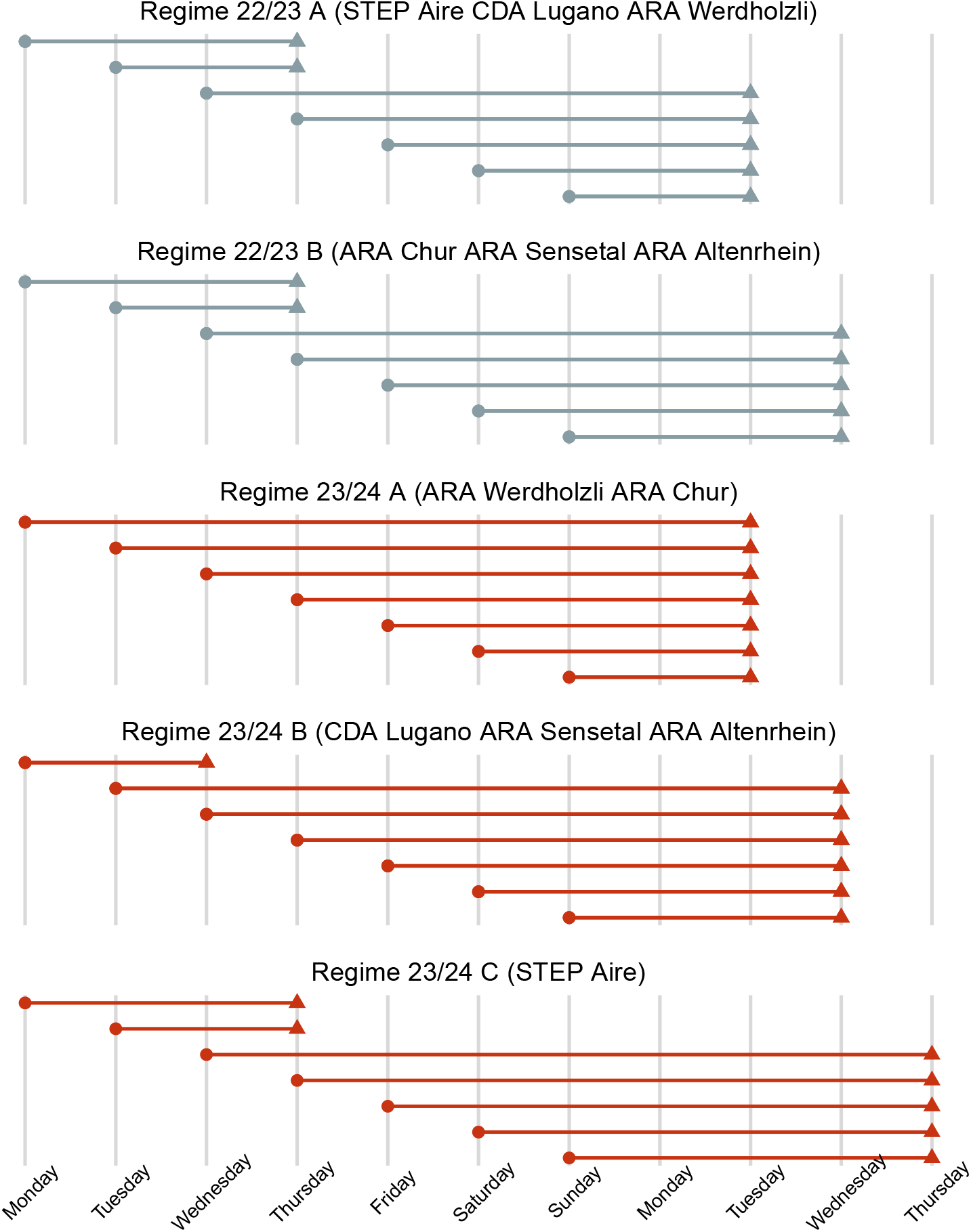
Regimes for collection and testing of wastewater samples by wastewater treatment plant. Circles show sample day and triangles show corresponding processing day. Color indicates the season where the regime was applied.

In addition to wastewater samples, each WWTP involved in our study provided measurements of the total daily inflow volume of wastewater, which we used to account for dilution of viral concentrations. More details on sample processing can be found in [18].

### 2.2 Time-varying populations estimated from mobile phone based mobility data

We partnered with mobility data consultancy *KIDO Dynamics* to generate estimates of time-varying population size in each WWTP catchment from call detail records (CDRs) of users of the *Sunrise* mobile phone network (35% market share in Switzerland). *KIDO Dynamics* generated estimates of anonymised mobile phone users’ movements within Switzerland and used these together with census data to estimate the time-varying population in each catchment. We extracted the time of arrival and duration of stay of individuals in each catchment on each day of the study period. From these extracts, we estimated the inferred population in each catchment in each hour of each day. To provide an indicative variable of time-varying population on a daily scale, for each day we calculated the mean number of users in each catchment per hour, calculated based on averaging the number of users for each hour, from here on referred to as “stays”.

### 2.3 Analysis of the viral concentration time-series

#### 2.3.1 Inference of viral degradation between collection and processing of wastewater samples

We assume that quantifiable viral concentrations *v*_*i*,*t*_ (where *i* indicates WWTP and *t* indicates the sample date) decay exponentially as a result of RNA degradation. Under this assumption, we developed a Bayesian Gamma regression model to estimate the rate of reduction of viral concentrations as a result of delay between collection and processing (*d*_*i*,*t*_) from the time series of measurements in 6 WWTPs, adjusting for the underlying national and local dynamics of viral concentrations. We calculated the delay between collection and processing as the number of days that elapsed between collection of the sewage sample and processing date according to the laboratory protocol (Figure 2). Furthermore, to account for changing dilution of RNA in the sample attributed to variations in the wastewater flow rates, we included the natural logarithm of the total daily influent flow (*f*_*i*,*t*_) at each WWTP as an additional fixed effect. We also correct for time-varying populations by including the natural logarithm of stays as a fixed effect, a time-varying population indicator (*m*_*i*,*t*_) as described in 2.2 (the log-transform confers the assumption that flow and stays have a linear relationship with RNA concentrations). Our model of viral concentration (in gene copies per liter [gc/L]) assumes a Gamma distribution with mean *µ*_*i*,*t*_ *>* 0 varying by WWTP *i* and over time *t*, and a common standard deviation *σ >* 0:

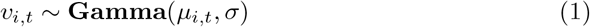

The mean, *μ*_*i*,*t*_, is expressed as:

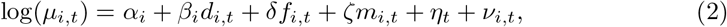

where *α*_*i*_ is a WWTP-specific intercept, *β*_*i*_, *δ*, and *ζ* are coefficients of fixed effects: testing delay (*d*_*i*,*t*_), log daily flow (*f*_*i*,*t*_), and log stays (*m*_*i*,*t*_), respectively. *η* and *ν*_*i*,*t*_ are national and local time-dependent components, respectively, capturing the underlying dynamics of prevalence in the population, and both were modelled as a first order random walk with penalised complexity priors [32]. The fixed effects were each assigned a Gaussian prior with mean of zero and a precision of 0.001.

We implemented the model using Integrated Nested Laplace Approximations as implemented in the `R-INLA R` package [33]. We allowed the delay effect to vary by WWTP (*β*_*i*_), the flow and stays effects (*δ*and *ζ*) were shared between all WWTPs. The general uncertainty in the gamma model is captured by modelling the standard deviation *σ*, which is shared between all sites and pathogens.

For comparison to previous estimates of degradation, we also present rates of degradation as the estimated time until 90% reduction in viral concentration measured (*T*_90_) [28, 30, 31]. This is calculated as

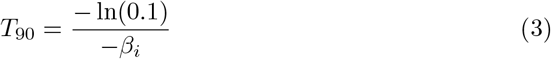

where *β*_*i*_ is the test delay effect as presented above. We report the *T*_90_ value only where we inferred a delay effect with 95% credible intervals that did not include zero.

#### 2.3.2 Quantifying the loss of measurement due to degradation

The limit of detection (LOD) and the limit of quantification (LOQ) are theoretical thresholds for viral concentration, which represent limitations in the digital PCR process to accurately detect and quantify viral RNA [17, 34]. We define the LOD as the threshold below which non-detection has a probability of greater than 5%. Previous analysis [34] of dPCR data suggests an LOD of 7.6 gc/ml per sample (LOD1), repeating the dPCR process for two replicates of the RNA extract reduce this to 3.8 gc/ml (LOD2). We define the limit of quantification as the concentration threshold below which the coefficient of variation is greater than 0.25. Previous studies have estimated this threshold to be 24 gc/ml [17, 34].

To establish the impact of RNA degradation on the quality of our concentration estimates, we quantified the number of measurements for which the quantifiable virus fell below the LOD and LOQ thresholds as a result of degradation between collection and processing. We then re-estimated the viral concentration on the day of sampling, using our model to predict the viral concentration with the delay set to zero days. We calculated the median number of measurements that were lost due to degradation by counting the days when the adjusted model predicts a value above each threshold with the 50% quantile of its posterior predictive distribution, while the actual measurement was below. We then estimated the lower and upper bounds of the 90% credible intervals as the number of measurements with the 95% quantile and 5% quantile of being above each threshold value, respectively. To exclude low-lying outliers unrelated to delay effects, we conditioned that 5% of the viral concentrations predicted by the adjusted model (delay effect removed) fell below each threshold. In our interpretation of our results, we focus on the LOD1 threshold, as this represents a concrete threshold where the risk of sample loss becomes unacceptable. We present results relative to the other thresholds alongside these results for the reader’s interest.

## Results

### 3.1 Inferred effect of processing delay on viral concentration measurements

In both seasons (October 2022 - March 2023 and October 2023 - March 2024), we found that the delay effect (*β*, corresponding to rate of exponential decay) varied substantially between pathogens and WWTPs (Table 1, Figure 3). Overall, the delay effect was largest for influenza A in both seasons, with mean effect ranging between −0.03 and −0.28 in 22/23 and −0.03 and −0.14 in 23/24 across WWTPs. There was a clear effect with delay (95% CrI not crossing 0) in Alternrhein, Chur, Sensetal and Werdhölzli in the 22/23 season and Chur, Sensetal and Lugano in the 23/24 season. These effects correspond to *T*_90_ value of between 8 and 23 days. The relative effects between WWTPs were similar for RSV (Figure 3), with the effects of −0.01 to −0.26 across both seasons. Sensetal was the highest with an effect of −0.26 (−0.36 − −0.15% 95% CrI). These rates correspond to a range of *T*_90_ values of 9 and 38 days among WWTPs with effects whose 95% CrI values did not cross 0. In the 22/23 season, the delay effect for influenza B and SARS-CoV-2, was generally lower with higher relative uncertainty such that the credible intervals bounded zero in most cases. The exceptions were Sensetal and Aire, where effects were measured for both pathogens, albeit smaller than those for IAV and RSV. There was a mean effect of −0.12 (−0.21 − −0.03 95% CrI) and −0.18 (−0.24 − −0.12 95% CrI) for IBV and SARS-CoV-2 respectively, in Sensetal, translating to mean *T*_90_ values of 19 and 12 days respectively. SARS-CoV-2 had a modest delay effect of −0.07 (−0.12 − −0.03 95% CrI) in Aire, which was similar to IVB −0.09 (−0.17 − −0.01 95% CrI), these result in *T*_90_ values of 31 and 25 days respectively. (Table 1). In the 23/24 season, effects remained unclear for IBV, with effect credible intervals crossing zero for both Aire and Sensetal, however a small effect was measured in Lugano −0.06 (−0.1 − −0.02 95% CrI). For SARS-CoV-2 however, the delay effect remained in Sensetal (−0.17 (−0.22 − −0.13 95% CrI)) alongside effects measured in Altenrhein (−0.10 (−0.16 − −0.03 95% CrI))and Chur (−0.08 (−0.12 − −0.03 95% CrI)) with *T*_90_ values of 13 to 31 days. The effect sizes were comparable between the seasons. This was most clear in the case of RSV where median values were most similar. However, the 95% credible interval ranges of delay effects overlapped between seasons in every WWTP for every pathogen. The inferred delay effect was largest in Sensetal for all pathogens and WWTPs in both seasons, with the exception of IBV in the 23/24 season. Conversely, the delay effect was most consistently low in Werdhölzli, although it was not necessarily the lowest in every scenario. In general the effects were larger for the 22/23 season for IAV and RSV however no such consistent pattern was clear for IBV and SARS-CoV-2.

**Fig. 3.**
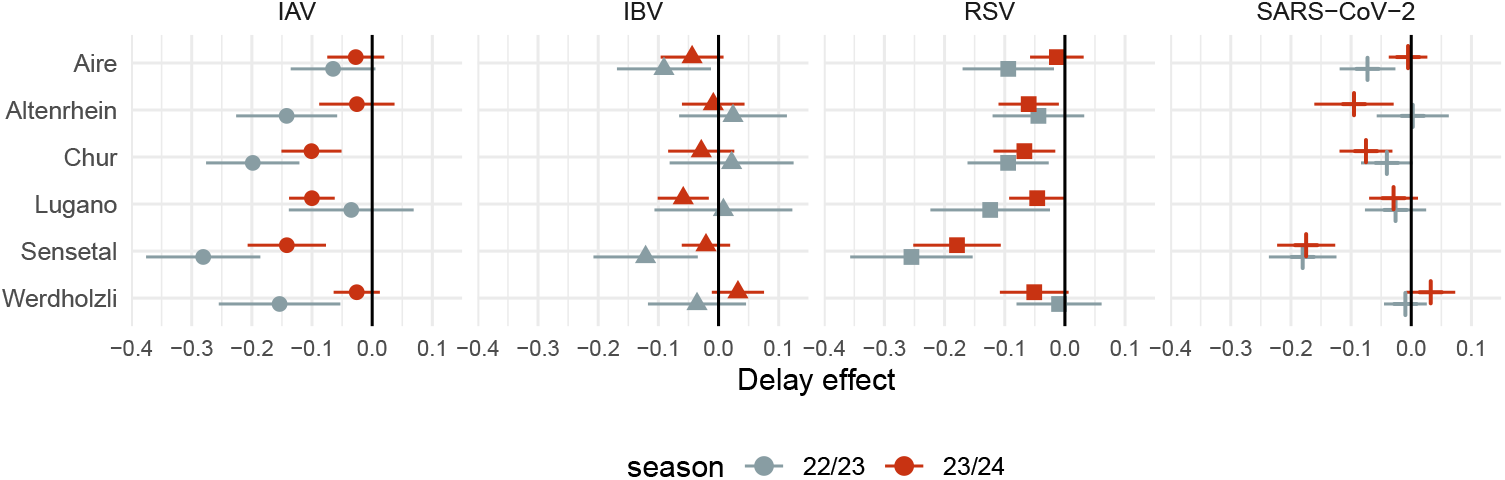
Posterior estimates of the coefficients of delay between collection and processing. Each panel shows size of delay effect for each pathogen, for each WWTP monitored. Color indicates the season where the effect was measured, error bars show the 95% quantiles of the marginal posterior distributions

**Table 1.**
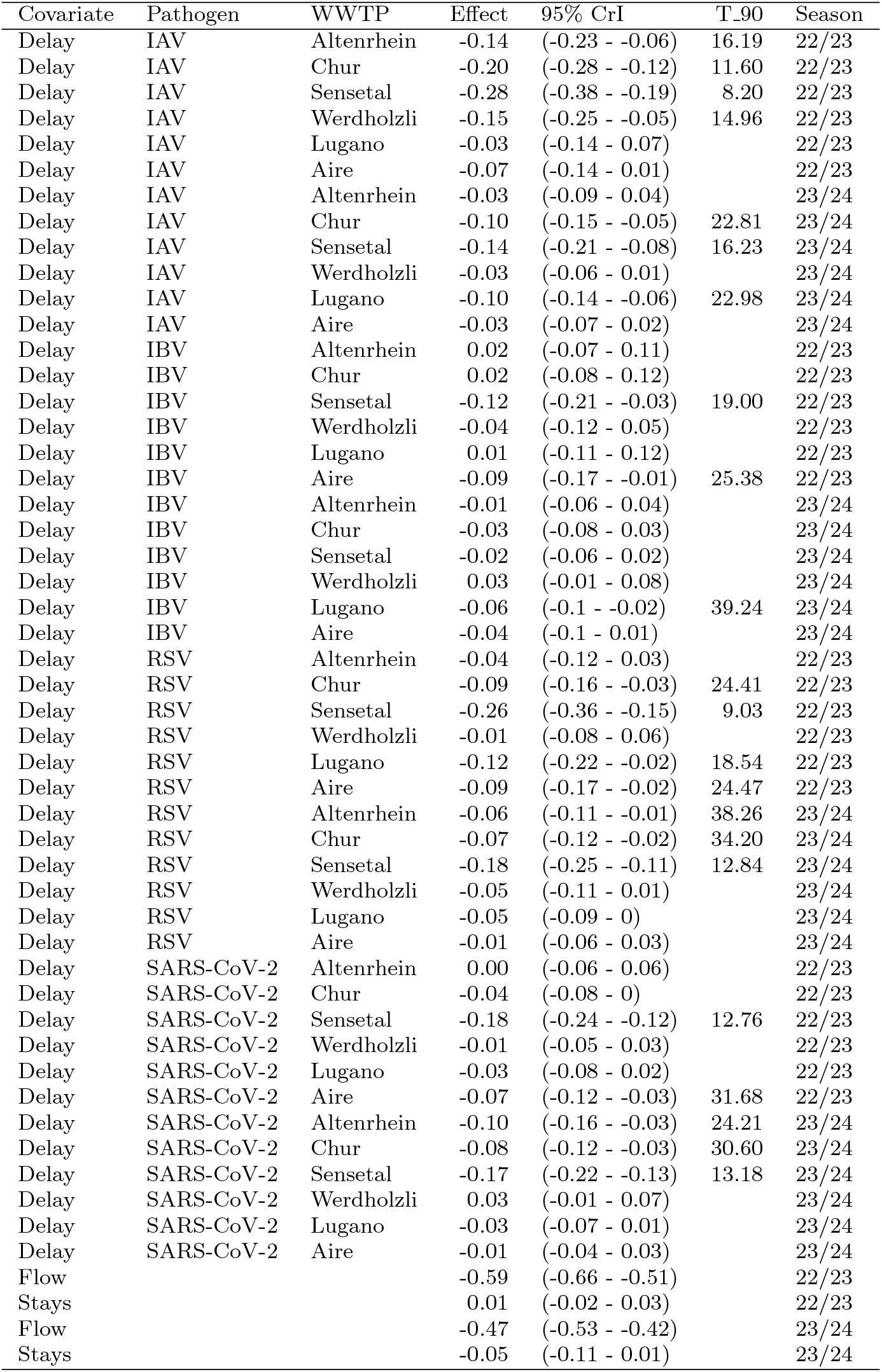
Effect of covariates onviral concentrations.

| Covariate | Pathogen | WWTP | Effect | 95% CrI | T_90 | Season |
| --- | --- | --- | --- | --- | --- | --- |
| Delay | IAV | Altenrhein | -0.14 | (-0.23 - -0.06) | 16.19 | 22/23 |
| Delay | IAV | Chur | -0.20 | (-0.28 - -0.12) | 11.60 | 22/23 |
| Delay | IAV | Sensetal | -0.28 | (-0.38 - -0.19) | 8.20 | 22/23 |
| Delay | IAV | Werdholzli | -0.15 | (-0.25 - -0.05) | 14.96 | 22/23 |
| Delay | IAV | Lugano | -0.03 | (-0.14 - 0.07) |  | 22/23 |
| Delay | IAV | Aire | -0.07 | (-0.14 - 0.01) |  | 22/23 |
| Delay | IAV | Altenrhein | -0.03 | (-0.09 - 0.04) |  | 23/24 |
| Delay | IAV | Chur | -0.10 | (-0.15 - -0.05) | 22.81 | 23/24 |
| Delay | IAV | Sensetal | -0.14 | (-0.21 - -0.08) | 16.23 | 23/24 |
| Delay | IAV | Werdholzli | -0.03 | (-0.06 - 0.01) |  | 23/24 |
| Delay | IAV | Lugano | -0.10 | (-0.14 - -0.06) | 22.98 | 23/24 |
| Delay | IAV | Aire | -0.03 | (-0.07 - 0.02) |  | 23/24 |
| Delay | IBV | Altenrhein | 0.02 | (-0.07 - 0.11) |  | 22/23 |
| Delay | IBV | Chur | 0.02 | (-0.08 - 0.12) |  | 22/23 |
| Delay | IBV | Sensetal | -0.12 | (-0.21 - -0.03) | 19.00 | 22/23 |
| Delay | IBV | Werdholzli | -0.04 | (-0.12 - 0.05) |  | 22/23 |
| Delay | IBV | Lugano | 0.01 | (-0.11 - 0.12) |  | 22/23 |
| Delay | IBV | Aire | -0.09 | (-0.17 - -0.01) | 25.38 | 22/23 |
| Delay | IBV | Altenrhein | -0.01 | (-0.06 - 0.04) |  | 23/24 |
| Delay | IBV | Chur | -0.03 | (-0.08 - 0.03) |  | 23/24 |
| Delay | IBV | Sensetal | -0.02 | (-0.06 - 0.02) |  | 23/24 |
| Delay | IBV | Werdholzli | 0.03 | (-0.01 - 0.08) |  | 23/24 |
| Delay | IBV | Lugano | -0.06 | (-0.1 - -0.02) | 39.24 | 23/24 |
| Delay | IBV | Aire | -0.04 | (-0.1 - 0.01) |  | 23/24 |
| Delay | RSV | Altenrhein | -0.04 | (-0.12 - 0.03) |  | 22/23 |
| Delay | RSV | Chur | -0.09 | (-0.16 - -0.03) | 24.41 | 22/23 |
| Delay | RSV | Sensetal | -0.26 | (-0.36 - -0.15) | 9.03 | 22/23 |
| Delay | RSV | Werdholzli | -0.01 | (-0.08 - 0.06) |  | 22/23 |
| Delay | RSV | Lugano | -0.12 | (-0.22 - -0.02) | 18.54 | 22/23 |
| Delay | RSV | Aire | -0.09 | (-0.17 - -0.02) | 24.47 | 22/23 |
| Delay | RSV | Altenrhein | -0.06 | (-0.11 - -0.01) | 38.26 | 23/24 |
| Delay | RSV | Chur | -0.07 | (-0.12 - -0.02) | 34.20 | 23/24 |
| Delay | RSV | Sensetal | -0.18 | (-0.25 - -0.11) | 12.84 | 23/24 |
| Delay | RSV | Werdholzli | -0.05 | (-0.11 - 0.01) |  | 23/24 |
| Delay | RSV | Lugano | -0.05 | (-0.09 - 0) |  | 23/24 |
| Delay | RSV | Aire | -0.01 | (-0.06 - 0.03) |  | 23/24 |
| Delay | SARS-CoV-2 | Altenrhein | 0.00 | (-0.06 - 0.06) |  | 22/23 |
| Delay | SARS-CoV-2 | Chur | -0.04 | (-0.08 - 0) |  | 22/23 |
| Delay | SARS-CoV-2 | Sensetal | -0.18 | (-0.24 - -0.12) | 12.76 | 22/23 |
| Delay | SARS-CoV-2 | Werdholzli | -0.01 | (-0.05 - 0.03) |  | 22/23 |
| Delay | SARS-CoV-2 | Lugano | -0.03 | (-0.08 - 0.02) |  | 22/23 |
| Delay | SARS-CoV-2 | Aire | -0.07 | (-0.12 - -0.03) | 31.68 | 22/23 |
| Delay | SARS-CoV-2 | Altenrhein | -0.10 | (-0.16 - -0.03) | 24.21 | 23/24 |
| Delay | SARS-CoV-2 | Chur | -0.08 | (-0.12 - -0.03) | 30.60 | 23/24 |
| Delay | SARS-CoV-2 | Sensetal | -0.17 | (-0.22 - -0.13) | 13.18 | 23/24 |
| Delay | SARS-CoV-2 | Werdholzli | 0.03 | (-0.01 - 0.07) |  | 23/24 |
| Delay | SARS-CoV-2 | Lugano | -0.03 | (-0.07 - 0.01) |  | 23/24 |
| Delay | SARS-CoV-2 | Aire | -0.01 | (-0.04 - 0.03) |  | 23/24 |
| Flow |  |  | -0.59 | (-0.66 - -0.51) |  | 22/23 |
| Stays |  |  | 0.01 | (-0.02 - 0.03) |  | 22/23 |
| Flow |  |  | -0.47 | (-0.53 - -0.42) |  | 23/24 |
| Stays |  |  | -0.05 | (-0.11 - 0.01) |  | 23/24 |

### 3.2 Quantifying the loss of measurements due to degradation

The WWTP at Sensetal had the highest proportion of measurements that we identified to have unacceptable loss rates due to sample degradation (falling below LOD1 - henceforth referred to as ‘lost’). Here we estimated a median of 33 (26 - 36 90% CrI) IAV measurements and 34 (30 - 40 90% CrI) RSV measurements in the 22/23 season and 25 (15 - 34 90% CrI) IAV measurements and 42 (28 - 48 90% CrI) RSV measurements in the 23/24 season were below the LOD1, however the delay effect adjusted predictions from our model indicated 50% chance of being above the LOD1 at the time of sampling (Table 2). We estimated that RSV and IAV had the most measurements lost to degradation across all WWTPs in the 22/23 season. Here IAV represented the greatest loss in four WWTPs (Altenrhein, Chur, Werdhölzli and Lugano), where RSV had the second highest estimated losses and had the highest losses in Aire and Sensetal and IAV had the second highest losses. All SARS-CoV-2 measurements were above the LOD1 during the 22/23 season (Table 2). We identified many more lost measurements of IBV in the 23/24 season, however viral concentrations were typically low and frequently bordering the LOD1 threshold and therefore many measurements below LOD1 were well within the predicted range by the delay adjusted model despite being identified as lost by our metric. Aside from IBV, we estimated that RSV and IAV lost a substantial number of measurements due to degradation between 6 and 39 for RSV and between 2 and 24 for IAV. We identified no lost measurements for SARS-CoV-2 in the 22/23 season and only in Sensetal in the 23/24 season, where an median estimate of 12 were identified as lost to degradation.

**Table 2.** Number of falling below LOQ and LOD as a result of reduction in concentrations due to post sample degradation.

| WWTP | Pathogen | Measurements lost to degradation |  |  |  |  |  |
| --- | --- | --- | --- | --- | --- | --- | --- |
|  |  | LOQ |  | LOD1 |  | LOD2 |  |
|  |  | median | 90% CrI | median | 90% CrI | median | 90% CrI |
| 22/23 Season |  |  |  |  |  |  |  |
| ARA Altenrhein | IAV | 2 | (0 - 4) | 15 | (8 - 20) | 7 | (5 - 7) |
| ARA Altenrhein | IBV | 0 | (0 - 0) | 1 | (0 - 3) | 12 | (0 - 23) |
| ARA Altenrhein | RSV | 1 | (0 - 1) | 2 | (0 - 4) | 7 | (1 - 12) |
| ARA Altenrhein | SARS-CoV-2 | 0 | (0 - 0) | 0 | (0 - 0) | 0 | (0 - 0) |
| ARA Chur | IAV | 2 | (2 - 5) | 14 | (8 - 19) | 8 | (6 - 8) |
| ARA Chur | IBV | 0 | (0 - 2) | 3 | (0 - 10) | 3 | (0 - 8) |
| ARA Chur | RSV | 0 | (0 - 0) | 5 | (2 - 7) | 5 | (1 - 6) |
| ARA Chur | SARS-CoV-2 | 0 | (0 - 0) | 0 | (0 - 0) | 0 | (0 - 0) |
| ARA Sensetal | IAV | 15 | (2 - 26) | 33 | (26 - 36) | 24 | (19 - 27) |
| ARA Sensetal | IBV | 2 | (0 - 3) | 3 | (3 - 7) | 17 | (3 - 27) |
| ARA Sensetal | RSV | 4 | (2 - 17) | 34 | (30 - 40) | 13 | (10 - 13) |
| ARA Sensetal | SARS-CoV-2 | 0 | (0 - 0) | 0 | (0 - 0) | 0 | (0 - 0) |
| ARA Werdholzli | IAV | 1 | (0 - 3) | 16 | (5 - 24) | 18 | (7 - 26) |
| ARA Werdholzli | IBV | 0 | (0 - 1) | 13 | (0 - 16) | 4 | (0 - 8) |
| ARA Werdholzli | RSV | 0 | (0 - 1) | 5 | (0 - 8) | 4 | (0 - 5) |
| ARA Werdholzli | SARS-CoV-2 | 0 | (0 - 0) | 0 | (0 - 0) | 0 | (0 - 0) |
| CDA Lugano | IAV | 0 | (0 - 1) | 9 | (0 - 16) | 5 | (0 - 8) |
| CDA Lugano | IBV | 0 | (0 - 0) | 3 | (0 - 6) | 3 | (0 - 15) |
| CDA Lugano | RSV | 1 | (1 - 2) | 11 | (4 - 20) | 7 | (5 - 11) |
| CDA Lugano | SARS-CoV-2 | 0 | (0 - 0) | 0 | (0 - 0) | 0 | (0 - 0) |
| STEP Aire | IAV | 1 | (0 - 2) | 5 | (2 - 14) | 4 | (1 - 8) |
| STEP Aire | IBV | 0 | (0 - 0) | 7 | (4 - 13) | 15 | (3 - 17) |
| STEP Aire | RSV | 0 | (0 - 0) | 12 | (6 - 17) | 18 | (9 - 26) |
| STEP Aire | SARS-CoV-2 | 0 | (0 - 0) | 0 | (0 - 0) | 0 | (0 - 0) |
| 23/24 Season |  |  |  |  |  |  |  |
| ARA Altenrhein | IAV | 3 | (0 - 3) | 2 | (0 - 15) | 21 | (1 - 22) |
| ARA Altenrhein | IBV | 0 | (0 - 0) | 7 | (0 - 37) | 10 | (0 - 13) |
| ARA Altenrhein | RSV | 3 | (1 - 4) | 21 | (5 - 25) | 10 | (6 - 13) |
| ARA Altenrhein | SARS-CoV-2 | 0 | (0 - 0) | 0 | (0 - 0) | 0 | (0 - 0) |
| ARA Chur | IAV | 1 | (1 - 1) | 9 | (5 - 16) | 7 | (5 - 7) |
| ARA Chur | IBV | 0 | (0 - 0) | 14 | (8 - 14) | 0 | (0 - 0) |
| ARA Chur | RSV | 2 | (0 - 6) | 9 | (5 - 12) | 3 | (3 - 3) |
| ARA Chur | SARS-CoV-2 | 0 | (0 - 0) | 0 | (0 - 0) | 0 | (0 - 0) |
| ARA Sensetal | IAV | 2 | (0 - 8) | 25 | (15 - 34) | 17 | (9 - 17) |
| ARA Sensetal | IBV | 0 | (0 - 0) | 0 | (0 - 37) | 17 | (8 - 23) |
| ARA Sensetal | RSV | 6 | (4 - 34) | 42 | (28 - 48) | 24 | (20 - 24) |
| ARA Sensetal | SARS-CoV-2 | 2 | (2 - 6) | 12 | (11 - 12) | 3 | (3 - 3) |
| ARA Werdholzli | IAV | 0 | (0 - 1) | 12 | (1 - 23) | 8 | (0 - 8) |
| ARA Werdholzli | IBV | 0 | (0 - 0) | 9 | (0 - 49) | 3 | (0 - 3) |
| ARA Werdholzli | RSV | 6 | (0 - 8) | 11 | (7 - 15) | 2 | (2 - 2) |
| ARA Werdholzli | SARS-CoV-2 | 0 | (0 - 0) | 0 | (0 - 0) | 0 | (0 - 0) |
| CDA Lugano | IAV | 0 | (0 - 0) | 15 | (10 - 26) | 11 | (10 - 11) |
| CDA Lugano | IBV | 0 | (0 - 3) | 18 | (10 - 21) | 1 | (1 - 1) |
| CDA Lugano | RSV | 4 | (2 - 5) | 14 | (2 - 21) | 4 | (3 - 4) |
| CDA Lugano | SARS-CoV-2 | 1 | (1 - 2) | 0 | (0 - 0) | 0 | (0 - 0) |
| STEP Aire | IAV | 0 | (0 - 1) | 8 | (2 - 15) | 2 | (1 - 2) |
| STEP Aire | IBV | 0 | (0 - 0) | 11 | (8 - 11) | 0 | (0 - 0) |
| STEP Aire | RSV | 1 | (0 - 5) | 6 | (0 - 14) | 0 | (0 - 0) |
| STEP Aire | SARS-CoV-2 | 0 | (0 - 0) | 0 | (0 - 0) | 0 | (0 - 0) |

In the case of Sensetal, the measurements identified as lost were consistently distributed throughout both seasons for influenza A and RSV, due to a combination of low overall concentrations and high degradation rates (Figure 4 A, Supplementary materials). However, for other WWTPs with high estimated measurement loss as a result of degradation, the majority of measurement loss occurred at the beginning and end of the seasonal wave, where concentrations were low. For example, for influenza A in Alternrhein, Chur, and Werdhölzli during the 22/23 season, where each was estimated to have had between 5 and 9 lost measurements within a window of less than two weeks at the beginning of the seasonal epidemic, followed by a long period of few lost measurements, before a similar period of high loss at the end of the season (Figure 4 B).

**Fig. 4.**
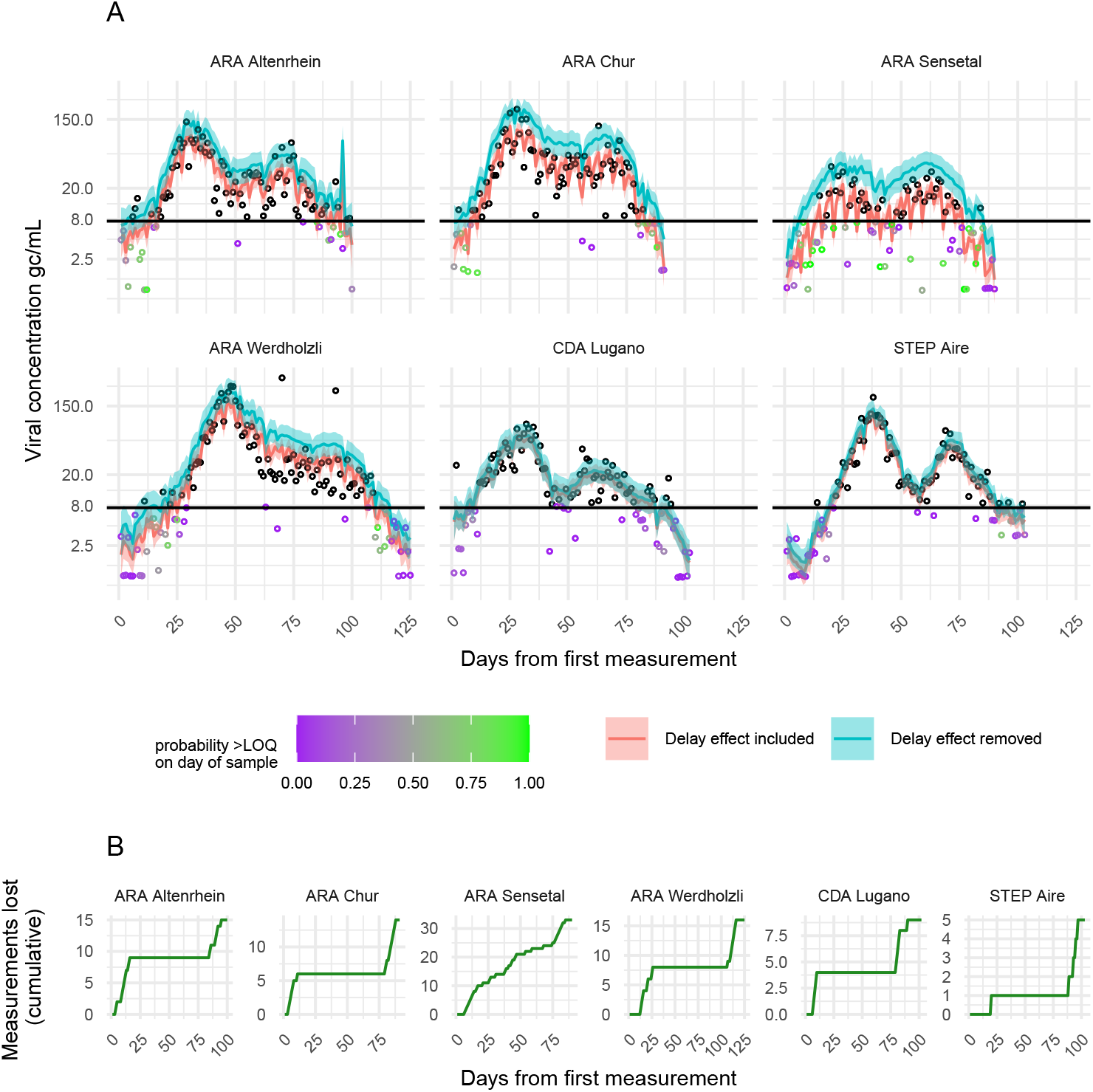
Effect of delay on modelled values of viral concentration of influenza A virus (IAV). A) Points show the viral concentrations measured on each day, the lines show the median modelled values, hue indicates whether the delay effect has been included or removed from the posterior predictions of viral concentration. The horizontal black line shows the theoretical limit of detection at 7.6 gc/mL. B) The cumulative number of measurements lost over the course of the season.

## 4 Discussion

Wastewater-based surveillance of respiratory infections relies on accurate quantification of viral concentrations in wastewater through PCR-based testing protocols. We used a Bayesian generalised linear model to quantify the rate at which the RNA of four respiratory pathogens (Influenza A and B, RSV and SARS-CoV-2) degraded in wastewater samples collected at 6 WWTPs. We found that, when accounting for overall temporal trends, delay between sample collection and processing was generally associated with a reduction in viral concentration. This reduction varied between pathogens and WWTPs. We found evidence that degradation between collection and processing of samples may be sufficient to impact the quality and reliability of viral concentration measurements. Reduction in viral concentrations has a particularly high impact on sample quality at the beginning of seasonal epidemics, which, in turn, may delay the detection of outbreaks.

Overall, our modelled outputs indicate that influenza A and RSV had the highest effect associated with the delay to testing. The delay effects inferred for SARS-CoV-2 or influenza B had 95% credible intervals that did not include 0 in only a subset of WWTPs. The consistency in our observations between two seasons under different testing protocols adds credence to the validity of our results.

Previous studies [30, 31] found varied evidence for SARS-CoV-2 degradation in samples stored at different temperatures. Ahmed et al. (2020)[31] estimated a *T*_90_ value of 27.8 (22.4 to 50.1 95% CrI) in untreated wastewater at 4^*o*^C. Although we were only able to detect significant delay effects for SARS-CoV-2 concentrations in two of our WWTPs during the 22/23 season and the three WWTPs during the 23/24 season, the effect measured in Aire in the 22/23 and Alternrhein and Chur in the 23/24 was close to this estimate with a *T*_90_ values of 32, 24 and 31 days respectively. The estimated effect in Sensetal was much stronger in both seasons with a *T*_90_ value of 12.9 (22/23 season) and 13.91 (23/24 season), which reflects much larger delay effects in this WWTP across all pathogens. Although Williams et al. (2024) [30] did not quantify the rate of degradation; they found that whether they observed degradation was dependent on the gene that was targeted by the PCR process. Our inability to sufficiently detect degradation in the remaining WWTPs may be due to lower rates of degradation being dominated by other sources of variability in concentrations and noise in measurements.

Nazir et al. (2010) [35] measured the rate of degradation of several strains of avian influenza A in distilled water, saline and surface water from Lake Constance at different temperatures. The highest rates of degradation were recorded in the surface water samples, although they did not study samples at 4^*o*^C they reported *T*_90_ values of 188 to 208 days for 0^*o*^C and 61 to 85 days for 10^*o*^C. This corresponds to lower rates of degradation than those we inferred for influenza A, however, this could be partially explained by differences in degradation rate in lake surface water and wastewater, since it has been found that a lower rate of SARS-CoV-2 degradation in tap water compared to untreated wastewater [31].

We quantified the implications of the degradation for wastewater monitoring applications by estimating the number of measurements that were below the limit of quantification due to RNA degradation between collection and processing. We found that Sensetal had the highest number of measurements identified as lost due to degradation. Here, a combination of a low viral concentration and a high degradation rate resulted in a large number of samples below the limit of detection (LOD1). Across all WWTPs, influenza A and RSV suffered the highest losses. In particular, for influenza A a large proportion of losses due to degradation occurred at the beginning and end of the influenza season. The frequent loss of data at the beginning of the season is important. One of the key benefits of wastewater-based surveillance and epidemiology is the potential for early detection and characterisation of outbreaks. Our results indicate that delays in testing could result in a longer period of measurements with insufficient concentrations to reliably quantify temporal trends in viral content. This could in turn delay timely epidemiological insights to support public health planning for the winter influenza season. The same phenomenon is present for RSV recorded in the 23/24 season (supplementary material). We expect we would observe a similar problem for RSV in the 22/23 season; however, regular testing for RSV began later in the RSV winter epidemic, when viral concentrations were already relatively high.

There was substantial variation in delay effect for the same pathogens in samples collected at different WWTPs in both seasons. The agreement between the variation in the delay effect between seasons under different testing protocols adds confidence that this variation is a true and systematic WWTP effect. The cause of this variation remains unclear, there are multiple factors that could contribute to this. For example, there is evidence that the content of the wastewater can substantially affect degradation of RNA. Most clearly, the pH value appears to have a large impact on viral concentrations [36]. There is evidence that ammonia content, presence of particular chemical compounds and overall viral concentration can impact the longevity of viral RNA [29]. This may correlate with differences in temporal variation in industrial activities that affect the composition of the wastewater at different times.

Our study represents the first attempt to quantify degradation of viral RNA between collection and processing of wastewater samples collected, stored and tested as part of a routine surveillance program. Our approach allows us to extend our observations of degradation to estimate the implications of degradation on data quality using metrics used in practice to monitor data quality. However, there are, a number of important limitations. Firstly, although our method allows variation in delay effects between pathogens and WWTPs, to support identifiabilty, we assume a constant rate of degradation over time. We anticipate that due to fairly consistent industrial enterprises in each catchment, the quality and content of the wastewater relevant to degradation is likely to vary more between WWTPs than over time within the WWTP; however, there may be some changes to the wastewater beyond those captured that substantially affect degradation of RNA differently at different times, for which we cannot adjust our model. Secondly, our model assumes that degradation occurs at a constant rate (exponential decay); evidence from clinical samples stored at lower temperatures shows that this rate may increase as overall concentrations decline [37]. This is likely to have a greater impact over longer degradation periods; however, since our delays do not exceed a few days, we propose that the approximation is reasonable. Thirdly, we assume that all samples are treated in the same way between collection and processing. In principle, this is true since all WWTPs follow a similar protocol; however, transit distance and time varies between each WWTP and the single laboratory where all samples are tested, and there may also be variation in the way WWTPs store the samples on the day of testing as the composite sample is accrued. Since this is not expected to vary systematically by day of the week, we do not believe that this is likely to affect our inferred delay effects; however, it may contribute to the variability between WWTPs through secondary factors. Although the samples are untreated, they have all been through some filtration before sampling occurs. The type of screening applied before sampling can vary between WWTPs, perhaps affecting their relative degradation rates when stored. We do not account for these differences and they may contribute to the between WWTP variation we observe. Additionally, because the concentrations of viral RNA are subject to multiple external factors, a certain degree of noise persists within the time series. This noise reduces the sensitivity of our model to detect lower rates of degradation. For this reason, we conclude that no clear reduction in viral concentrations is observed in many instances where some degradation may be present at rates too low to reliably detect in the context of our study. Finally, the limits of detection and quantification used in our analysis only include the variability introduced during the digital PCR process [34]. Additional sources of variability in detection may impact the detectability and quantification of viral RNA in wastewater. However, based on our own analyses we anticipate this variation to be small compared to those involved in the dPCR process.

## 5 Conclusions

Using a Bayesian time series model of viral concentrations in wastewater collected as part of a national surveillance program, we were able to identify that the delay between collection and processing was associated with a reduction in viral concentration. The effect of this delay broadly agreed with previous estimates from laboratory tests. We found that degradation appears to be more pronounced in influenza A and RSV than in SARS-CoV-2 and influenza B. There was substantial variability in the strength of degradation among WWTPs. The degradation we measured has the most meaningful impact for Wastewater-based influenza A surveillance, where we inferred that a large proportion of samples collected during the early period of the season were below the limit of quantification due to degradation. We advocate for further evaluation of the variation in degradation rates between WWTPs to establish whether particular sampling protocols can improve the preservation of RNA in samples for longer. We propose that focused sampling and testing protocols can improve the quality of data during these key periods of the seasonal epidemic, where viral concentrations remain relatively low.

## Acknowledgments

The authors wish to thank KIDO dynamics for their support of in the preparation and provision of the mobile phone based mobility data used in this study, in particular Alberto Hernando, David Mateo and Christian Garcia. We also wish to thank the many personnel involved between our partner treatment plants and at Eawag involved in the testing and processing of wastewater samples.

## Funding

JDM, JDK, CG, CO, TRJ and TS received funding from the Swiss National Science Foundation grant number 205933. AL and JR received no funding for this project.

### Data and code availability

Data and code are available at https://github.com/wise-ch/wwperiodicity. Mobility data available by request from KIDO Dynamics

### Author contribution

The work was conceived by JDM and JKE. Data curation was performed by JDM, JKE, CG and AL. The methodology and formal analysis were performed by JDM. Supervision was provided by JR, CO, TRJ and TS. The manuscript was written by JDM. The manuscript was reviewed and edited by all of the authors.

