## Supplementary material for "Post-sampling degradation of viral RNA in wastewater impacts the quality of PCR-based concentration estimates"

James D. Munday

June 6, 2025

### **1 Supplemental figures**

#### **1.1 Flow and mobility timeseries**

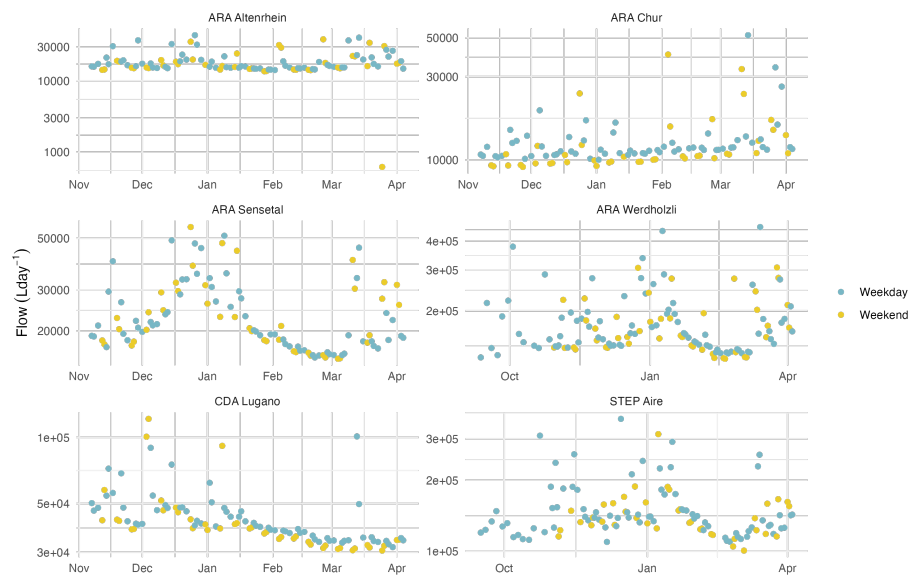

Figure 1: Influent wastewater per day by treatment plant 22/23

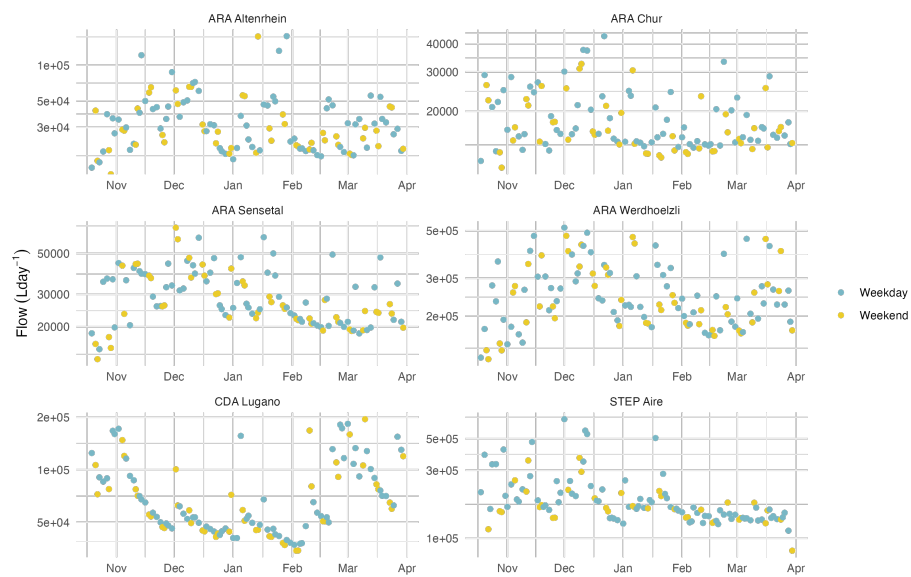

Figure 2: Influent wastewater per day by treatment plant 23/24

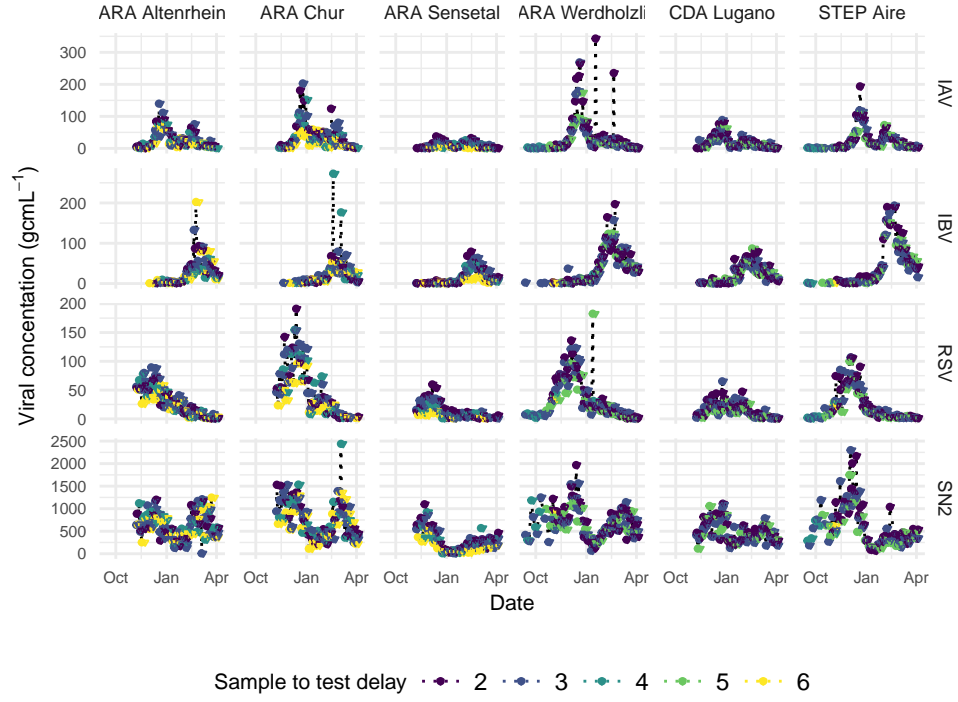

Figure 3: Time-series of viral load for four pathogens (top to bottom) in 6 WWTPs (left to right) color of points shows the delay between collection and testing (data shown for October 2022 - aMarch 2023)

### 1.2 Flow and mobility timeseries

We applied Welch’s Periodogram algorithm to establish the frequencies present in the viral loads. We found that in all WWTPs there was elevated contribution from periods in the order of 1 week (Figure 3). In particular, periodicity was strongest for RSV and Influenza viruses. SARS-CoV-2 viral concentrations showed lower intensity of periodicity at seven days, but still displayed increased PSD values between 5 and 8 day period. The WWTP of Sensetal in canton Bern also had a clearer peak at 7 days than the other WWTPs for RSV, IAV and IBV. In particular IAV showed much clearer periodicity in Sensetal than the other WWTPs, where the periodicity was comparable to SARS-CoV-2.

### 1.3 Posterior predictive plots

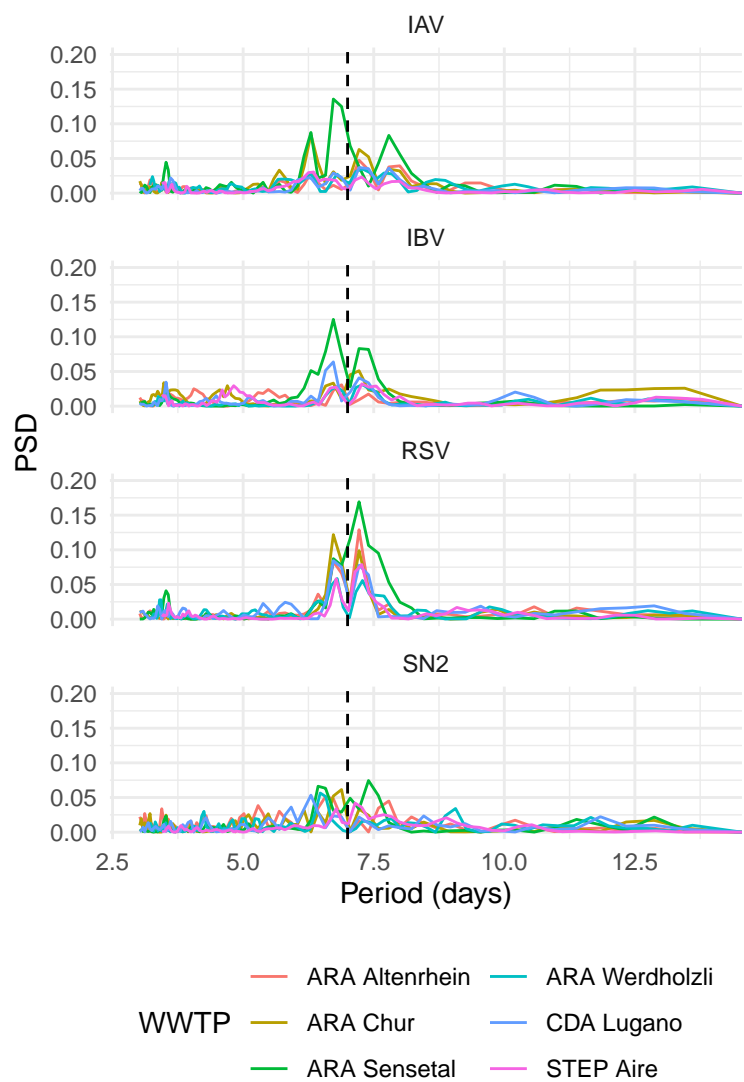

Figure 4: Power spectral density of viral load time-series of each pathogen at each WWTP

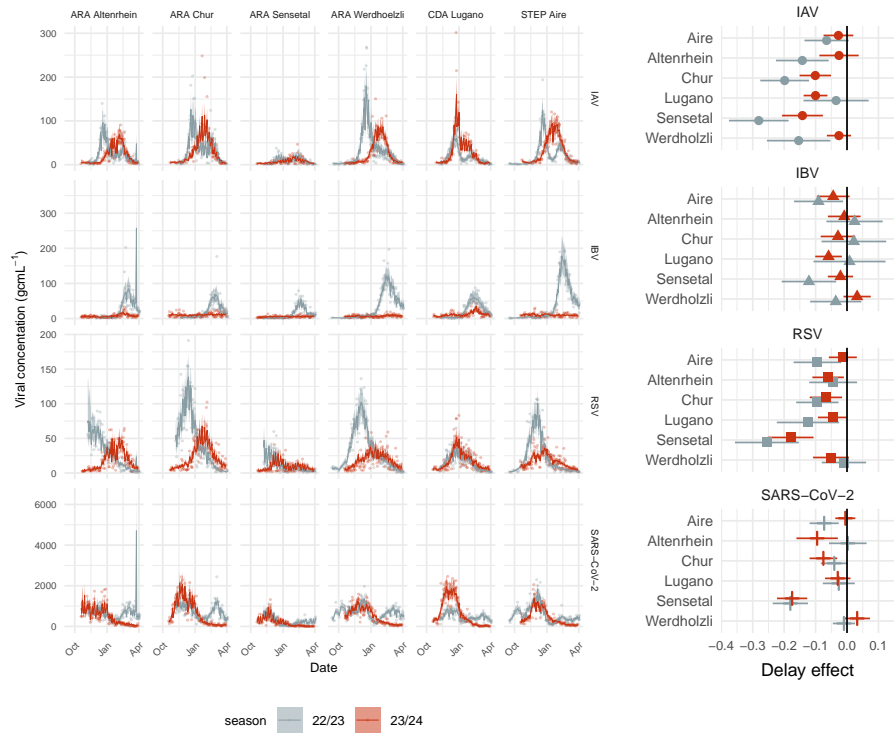

Figure 5: Posterior predictive plot of the time-series model for viral concentrations in each season

### 1.4 Measurements lost to degradation

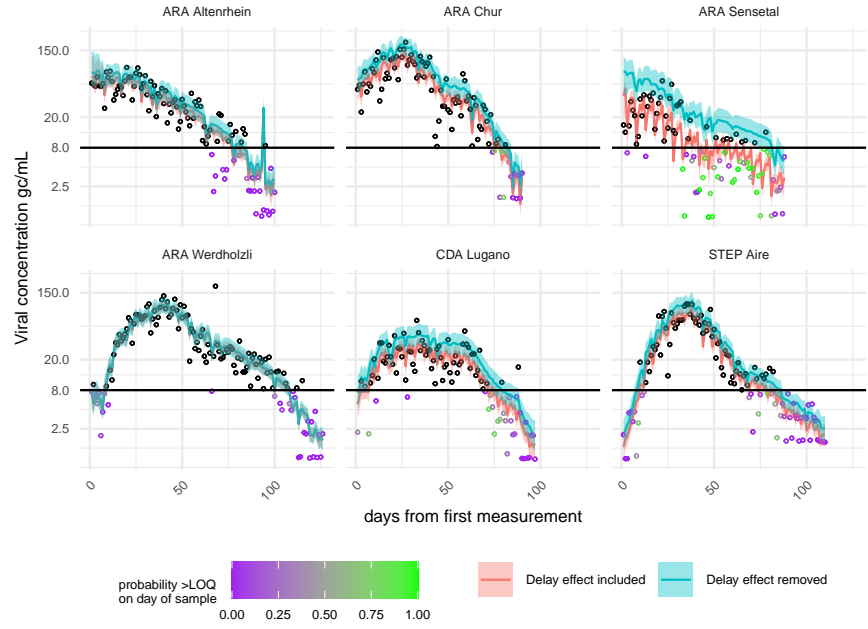

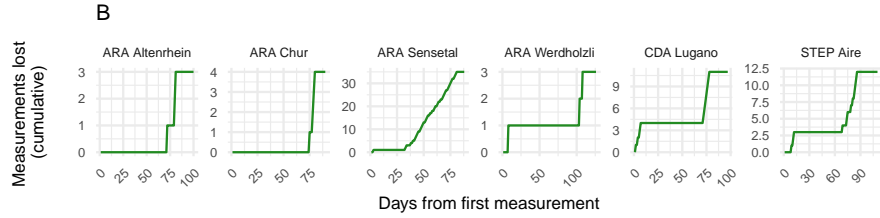

Figure 6: Effect of delay on modelled values of viral concentration of Respiratory Sincycal Virus (RSV) during the 22/23 season. Points show the viral concentrations measured on each day, the lines show the median modelled values, hue indicates whether the delay effect has been included or removed from the posterior predictions of viral concentration. The horizontal black line shows the theoretical limit of quantification at 8 gc/mL.

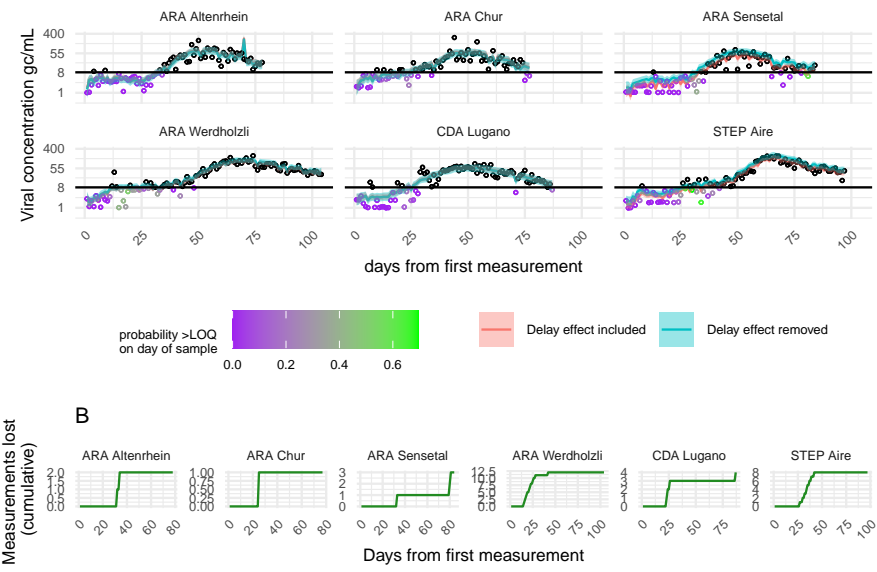

Figure 7: Effect of delay on modelled values of viral concentration of influenza B virus (IBV) during the 22/23 season. Points show the viral concentrations measured on each day, the lines show the median modelled values, hue indicates whether the delay effect has been included or removed from the posterior predictions of viral concentration. The horizontal black line shows the theoretical limit of quantification at 8 gc/mL.

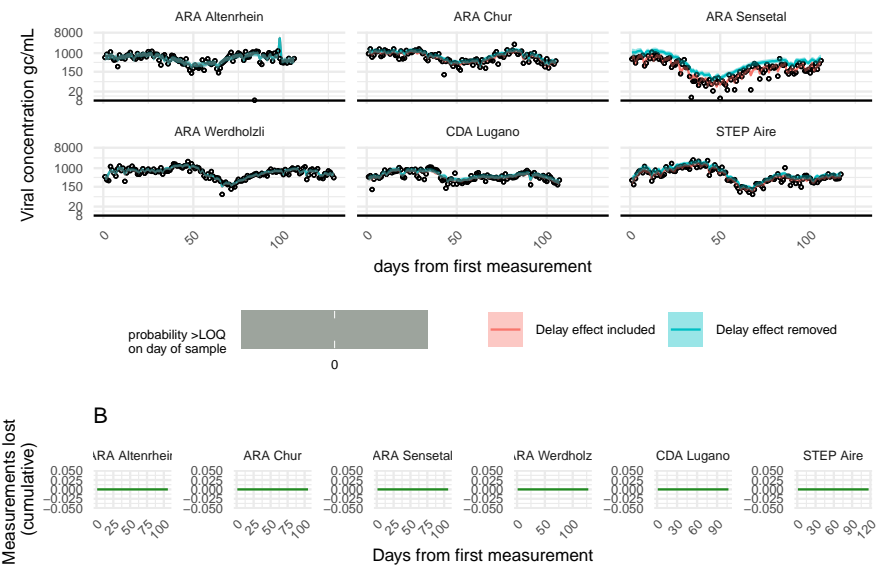

Figure 8: Effect of delay on modelled values of viral concentration of SARS-CoV-2 during the 22/23 season. Points show the viral concentrations measured on each day, the lines show the median modelled values, hue indicates whether the delay effect has been included or removed from the posterior predictions of viral concentration. The horizontal black line shows the theoretical limit of quantification at 8 gc/mL.

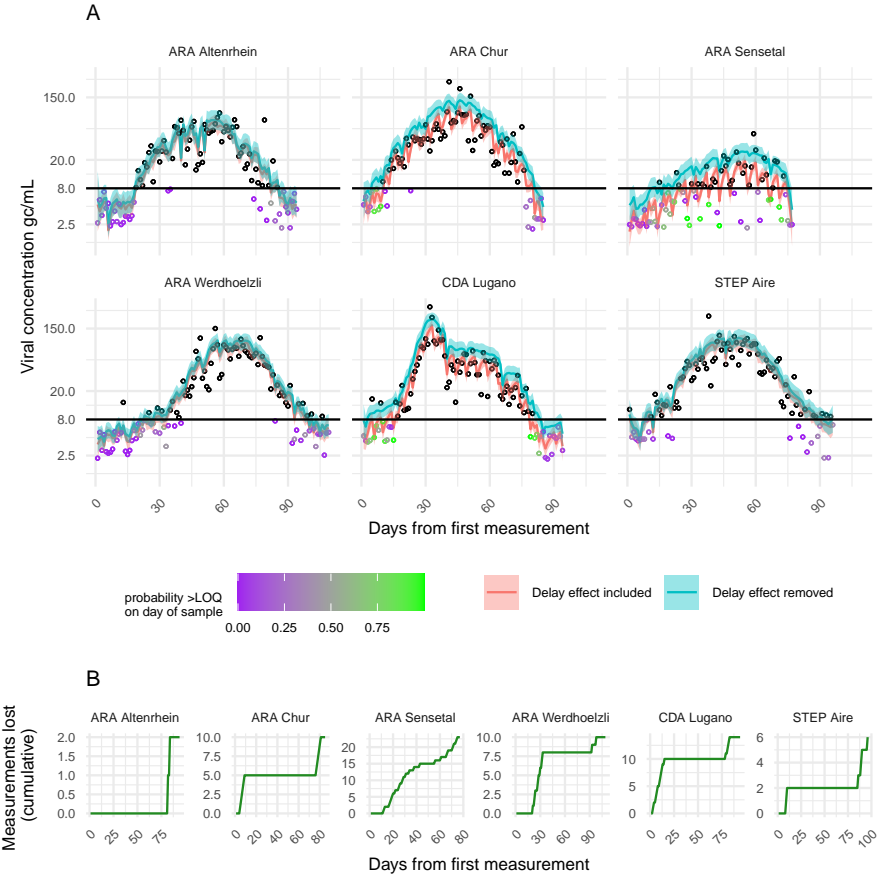

Figure 9: Effect of delay on modelled values of viral concentration of influenza A virus (IAV) during the 23/24 season. Points show the viral concentrations measured on each day, the lines show the median modelled values, hue indicates whether the delay effect has been included or removed from the posterior predictions of viral concentration. The horizontal black line shows the theoretical limit of quantification at 8 gc/mL.

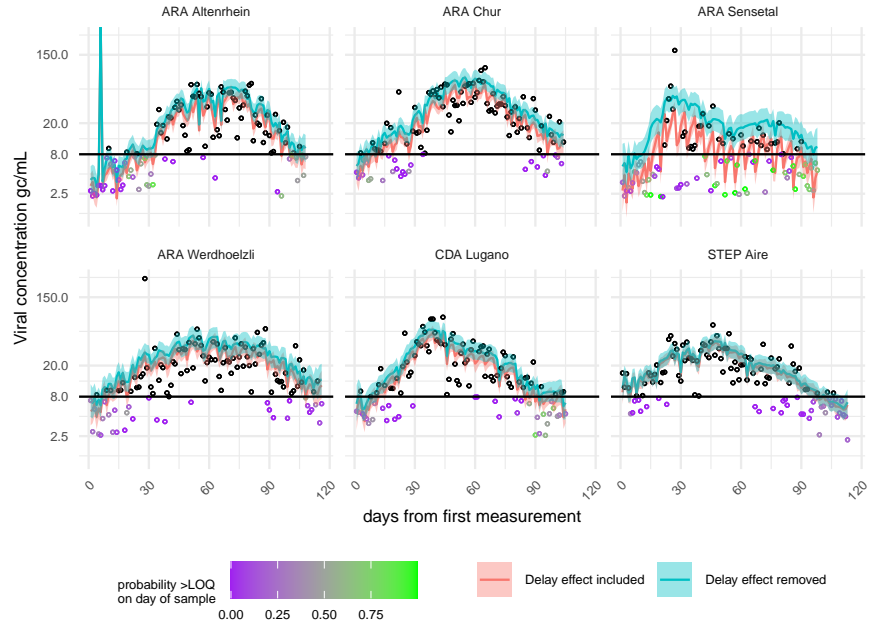

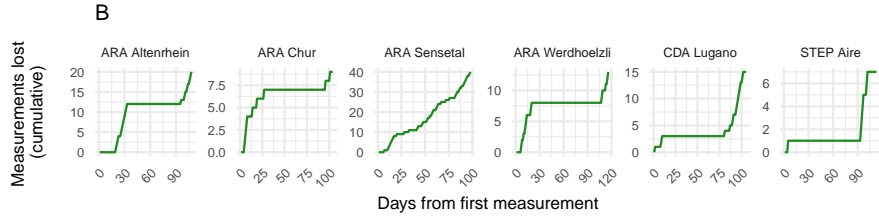

Figure 10: Effect of delay on modelled values of viral concentration of Respiratory Sincycal Virus (RSV) during the 23/24 season. Points show the viral concentrations measured on each day, the lines show the median modelled values, hue indicates whether the delay effect has been included or removed from the posterior predictions of viral concentration. The horizontal black line shows the theoretical limit of quantification at 8 gc/mL.

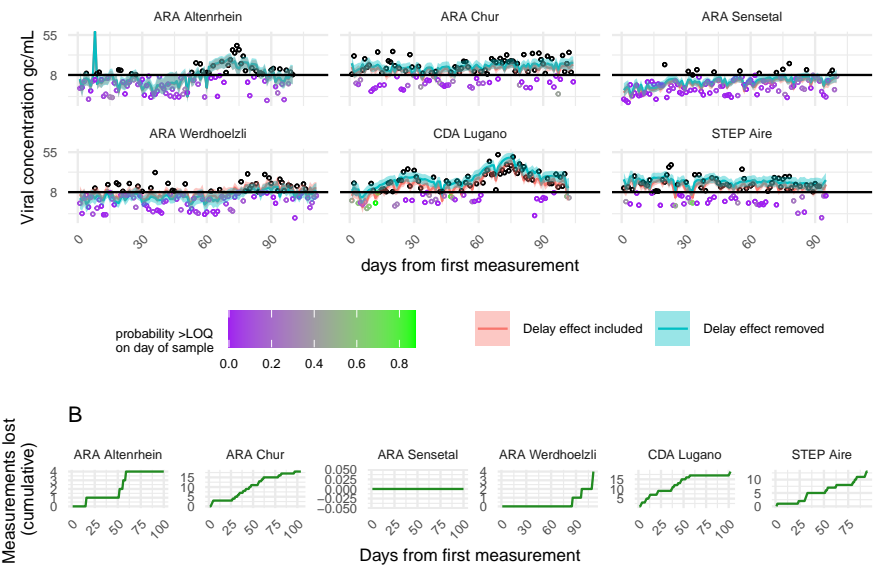

Figure 11: Effect of delay on modelled values of viral concentration of Influenza B virus (IBV) during the 23/24 season. Points show the viral concentrations measured on each day, the lines show the median modelled values, hue indicates whether the delay effect has been included or removed from the posterior predictions of viral concentration. The horizontal black line shows the theoretical limit of quantification at 8 gc/mL.

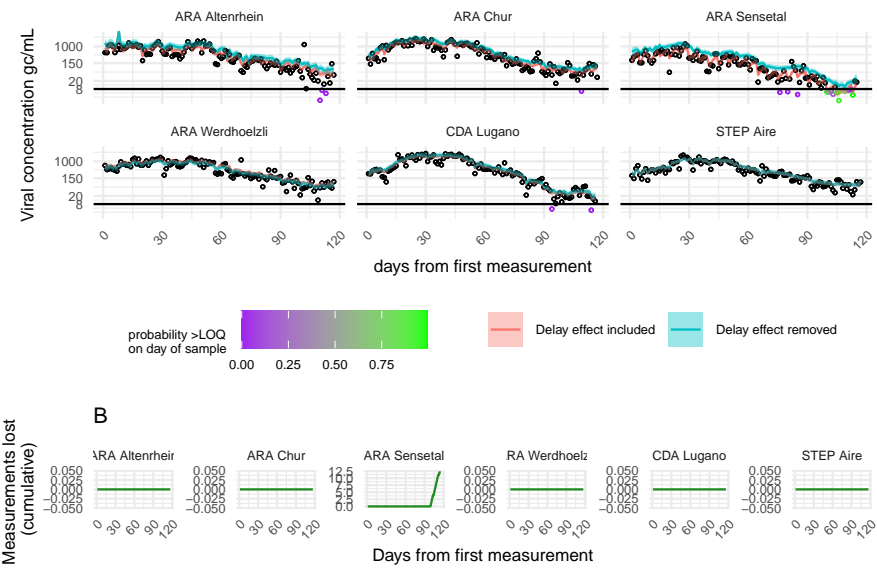

Figure 12: Effect of delay on modelled values of viral concentration of SARS-CoV-2 during the 23/24 season. Points show the viral concentrations measured on each day, the lines show the median modelled values, hue indicates whether the delay effect has been included or removed from the posterior predictions of viral concentration. The horizontal black line shows the theoretical limit of quantification at 8 gc/mL.
